# Dissociable roles for the striatal cholinergic system in different flexibility contexts

**DOI:** 10.1101/2021.12.22.473508

**Authors:** Brendan Williams, Anastasia Christakou

## Abstract

The production of behavioural flexibility requires the coordination and integration of information from across the brain, by the dorsal striatum. In particular, the striatal cholinergic system is thought to be important for the modulation of striatal activity. Research from animal literature has shown that chemical inactivation of the dorsal striatum leads to impairments in reversal learning. Furthermore, proton magnetic resonance spectroscopy work has shown that the striatal cholinergic system is also important for reversal learning in humans. Here, we aim to assess whether the state of the dorsal striatal cholinergic system at rest is related to flexible behaviour in reversal learning. We provide preliminary results showing that variability in choline in the dorsal striatum is significantly related to both the number perseverative and regressive errors that participants make, and their rate of learning from positive and negative prediction errors. These findings, in line with previous work, suggest the resting state of dorsal striatal cholinergic system has important implications for producing flexible behaviour. However, these results also suggest the system may have heterogeneous functionality across different types of tasks measuring behavioural flexibility. These findings provide a starting point for further interrogation into understanding the functional role of the striatal cholinergic system in flexibility.

## Introduction

Behavioural flexibility enables an individual to generate complex behaviour that enables them to adaptively respond changes in their world. The neurotransmitter acetylcholine is thought to play a critical role in this ability (Yamanaka et al., 2018). Evidence of the importance of the striatal cholinergic system has mostly come from the animal literature, where cholinergic interneurons in the dorsomedial striatum have been shown to be important for flexibility (Bradfield & Balleine, 2017). The reversal learning task is a commonly used paradigm for studying behavioural flexibility, with reversal learning associated with increases in acetylcholine release in the dorsomedial striatum (Ragozzino et al., 2009). Cholinergic neurotransmission modulates medium spiny neuron activity directly via the expression of muscarinic receptors on medium spiny neurons (Assous, 2021), and indirectly via the expression of acetylcholine receptors on glutamatergic and dopaminergic projection neurons, GABAergic interneurons, and as autoreceptors on cholinergic interneurons (Ding et al., 2010; English et al., 2012; Kljakic et al., 2017; Kreitzer, 2009). Inactivation of cholinergic interneurons, and antagonism of cholinergic receptors on medium spiny neurons in the dorsomedial striatum impairs reversal learning performance (McCool et al., 2008; Ragozzino et al., 2009; Tzavos et al., 2004). These impairments, as indexed by a reduced ability to update outcome contingencies following reversal and an increase in regressive errors are seen following the loss of cholinergic interneuron activity, or input from parafascicular nucleus of the thalamus to cholinergic interneurons (Brown et al., 2010; Ragozzino et al., 2002).

The response of cholinergic interneurons to changes in outcome contingency is thought to be dependent on input from the parafascicular nucleus of the thalamus in rodents, and its homologue in human and non-human primates, the centromedian-parafascicular nuclei (Smith et al., 2011). Unlike other thalamostriatal pathways, connections between the centromedian-parafascicular nuclei and the striatum show preferential connectivity with cholinergic interneurons in the striatum (Smith et al., 2009). Moreover, these thalamostriatal connections are crucial for the role of cholinergic interneurons for flexible behaviour. For instance, inactivation of parafascicular nucleus in rodents impaired reversal learning performance, with similar behavioural impairments seen in studies where cholinergic interneurons were chemically inactivated (Brown et al., 2010). The importance of input from the parafascicular nucleus to the striatal cholinergic system is thought to be specific for reversal learning, and direct evidence for the role of these thalamostriatal connections for flexible behaviour is provided by Bradfield et al. (2013). Firstly, they show bilateral lesions of the parafascicular nucleus impair reversal, but not initial learning. Next contralateral lesions of the parafascicular nucleus and dorsomedial striatum, but not ipsilateral lesions of the same regions are also shown to impair reversal learning. This is because unilateral lesions spare thalamostriatal connections in the hemisphere contralateral to the lesions, while contralateral lesions leave no intact connectivity because at least one node of the circuit is ablated in each hemisphere. Finally, unilateral lesions of the parafascicular and the chemical inactivation of cholinergic interneurons in the contralateral hemisphere also leave reversal learning impaired, emphasising that it is the inactivation of these thalamic connections specifically that impair reversal learning, providing compelling evidence for the role thalamostriatal connectivity with the striatal cholinergic system in the production of behavioural flexibility.

Compared with animal research, where invasive experiments can interrogate causal interactions, studying the role of thalamostriatal connectivity and the striatal cholinergic system in behavioural flexibility in humans is not trivial. Nevertheless, proton magnetic resonance spectroscopy (^1^H-MRS) is a non-invasive application of nuclear magnetic resonance spectroscopy, and is used to measure brain metabolites *in vivo* (Keeler, 2010). Theoretically, ^1^H-MRS can be used to directly measure acetylcholine, but its concentration is so low *in vivo* that its signal is masked by other choline containing compounds (Bell, Lindner, et al., 2019). These choline containing compounds are choline, glycerophosphocholine (GPC), and phosphocholine (PC). These metabolites could be used to indirectly study acetylcholine function. For instance, choline is the rate-limiting factor in synthesis of acetylcholine (Lockman & Allen, 2002), and synaptic choline levels are related to cholinergic interneuron activity with prolonged activation of cholinergic interneurons decreasing the concentration of choline in the synaptic cleft (Löffelholz, 1998). However, typical ^1^H-MRS approaches to quantifying choline containing compounds model them as a single peak due to their proximity on the spectrum. Doing so masks any functionally relevant choline effects as choline concentrations are anti-correlated with other choline-containing compounds (Lindner et al., 2017). Therefore, if we were able to separably measure choline from GPC and PC, then we could use this to indirectly and non-invasively study cholinergic system in humans. Previous work from our lab has demonstrated that choline can be separated from GPC and PC using ^1^H-MRS at three tesla by modelling choline as a separate peak from a combined GPC and PC peak. We observed task related functional changes in choline that were in line with expected changes in acetylcholine release during visuospatial attention (Lindner et al., 2017) which suggests that quantifying choline separately from GPC+PC using ^1^H-MRS may be an appropriate proxy for measuring acetylcholine activity in vivo.

We have previously used ^1^H-MRS to study the role of the dorsal striatal cholinergic system in behavioural flexibility using a multi-alternative probabilistic reversal learning task. Functional ^1^H-MRS was previously used by Bell et al. (2018) to study changes in choline that functionally relate to flexible behaviour. ^1^H-MRS data were acquired in the dorsal striatum while participants completed a multi-alternative probabilistic reversal learning task, and levels of choline and GPC+PC were quantified from metabolite spectra. The reversal of reward contingencies coincided with a significant decrease in the concentration of choline, but not GPC+PC or the total sum of choline containing metabolites, in line with previous findings of choline kinetics following the stimulation of cholinergic neurons in animals (Löffelholz, 1998), and from previous work using ^1^H-MRS to study visuospatial attention (Lindner et al., 2017). These results show the functional relevance of choline for behavioural flexibility and demonstrate the specificity of this metabolite as a proxy for acetylcholine release.

Performance during reversal learning can be summarised in several ways. Direct measures of performance include the number of trials taken to reach a predefined learning criterion, number of correct responses, or the number of perseverative and regressive errors participants make. Following the reversal of reward contingencies in the task, the continued selection of the previously correct response strategy is known as response perseveration and the number trials before switching to using a difference response strategy is a measure of perseverative errors. Following a change in response strategy and in the absence of any reversal of outcome contingencies, trials where participants revert to using the now incorrect response strategy are used to measure regressive errors. Latent variables of performance can be inferred by fitting models to behavioural data. For instance, temporal difference reinforcement learning models can be used to model how participants learn from experience (Sutton & Barto, 2018). These models describe how people learn associations between actions and outcomes. Learning is driven by reward prediction errors, which describe the difference between actual and expected outcomes and are used to generate future estimates of expected value (Schultz et al., 1997). The rate of expected value updating is determined by the learning rate, and can be symmetric (a single learning rate α) or asymmetric for positive (α^+^) and negative (α^-^) prediction errors (Niv et al., 2012). Reversal learning performance is associated with dorsal striatal choline levels at rest, with Bell, Lindner et al. (2019) finding choline concentrations were positively correlated with perseverative errors, and negatively correlated with α^-^. Additionally, α^-^ was negatively correlated with perseverative errors during reversal learning. These results show that lower levels of choline in the dorsal striatum at rest are associated with a quicker change in behaviour following the onset of reversal during multi-alternative probabilistic reversal learning and suggests that participants who reversed more quickly had lower levels of acetylcholine at rest, or more efficient re-uptake of choline following acetylcholine release.

In comparison to multi-alternative probabilistic reversal learning, two-choice serial reversal learning task is computationally simpler to solve. This simplicity means participants can feasibly complete multiple reversals over the course of the task. In Williams & Christakou, (2021) we show that functional connectivity between the centromedian-parafascicular nuclei and the associative dorsal striatum is significantly increased during the processing of negative feedback relative to positive feedback. This change in functional connectivity could reflect a general error signal from the thalamus to cholinergic interneurons to promote flexible behaviour. However, these results do not directly implicate the striatal cholinergic system in the generation of flexible behaviour. Therefore, we next want to use ^1^H-MRS to determine whether, in line with animal literature and our previous human work, serial reversal learning performance is associated with the striatal cholinergic system. More specifically, we are interested in the relationship between reversal learning and choline in the dorsal striatum at rest. Participants completed a probabilistic reversal learning task and we then acquired spectroscopy data from the dorsal striatum while at rest. Based on previous results, we predict reversal learning performance, as indexed by perseverative and regressive errors and parameter estimates from reinforcement learning models, will be associated with levels of choline in the dorsal striatum at rest.

## Methods

### Participants

Thirty three healthy adult participants were recruited to take part in this study. Thirty one of these participants were a subset of participants who also took part in the study described in Williams & Christakou, (2021). Participants were recruited through opportune sampling within the University of Reading community. Eligible participants were right-handed, and self-reported no use of cigarettes, recreational drugs, prescription of psychoactive medication, and that they had no formal diagnosis of a psychiatric or neurological condition. Participants received £15 compensation for their time. Participants were included in the analysis reported here if they responded on at least 95% of the trials in the learning task, if their MRS spectral acquisition appeared correctly aligned within the striatum on their T1 acquisition, and if we were able to quantify separate peaks for choline and glycerophosphocholine plus phosphocholine. Three participants were excluded because they responded on fewer than 95% of trials, one had registration issues, one had no behavioural data, one was manually removed as their behaviour suggested they did not understand the task, one had spectroscopy data lost, four had spectra that were corrupted during acquisition, and nine had choline peaks that could not be separated. Our sample used for statistical analyses consisted of thirteen participants (mean age = 22.69 years; SD = 3.20; range = 18-29; 11 female). The study was approved by the research ethics committee of the University of Reading [UREC 19/42].

### Probabilistic reversal learning task

This task has been described previously in detail in Williams & Christakou, (2021). Two abstract images of fractal patterns were shown on the left and right hemifield of the visual display. Participants had to choose one of the two images within 2000ms by pressing the corresponding button on a button box, else a “ too late” message was displayed. The outcome of the participant’s choice was then presented, followed by their cumulative points total. Figure 1 shows a schematic of the task trial structure and timings.

**Figure 1.**
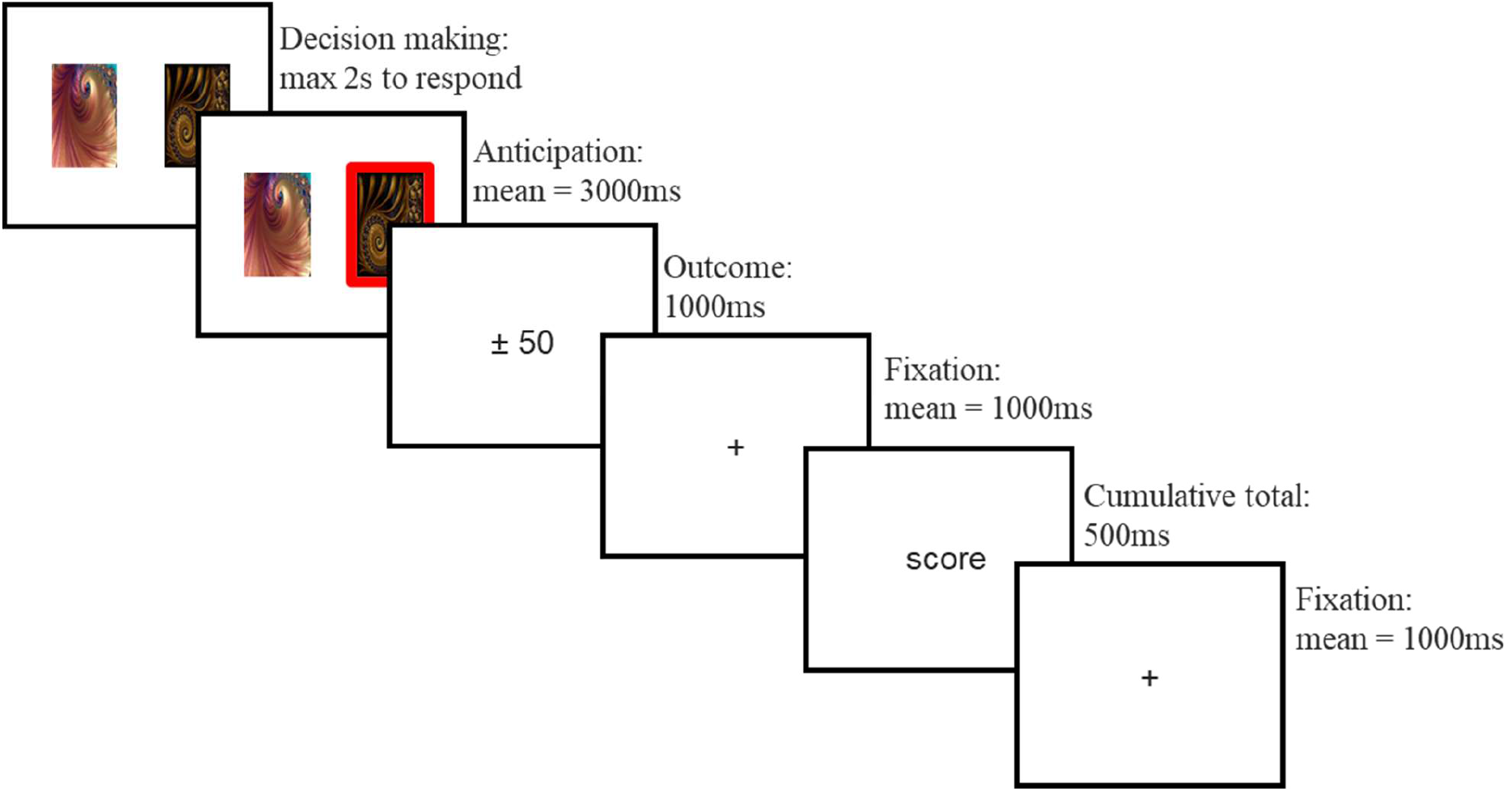
Overview of a single trial. Participants are initially shown two abstract fractal images and given two seconds to choose one image. Their choice is then highlighted. The participant is then shown the outcome of their choice; this will either be an increase or decrease of 50 points if they selected an image, or 0 points if they made no choice. The outcome is followed by a fixation cross, their cumulative total so far, and finally another fixation cross.

**Figure 2.**
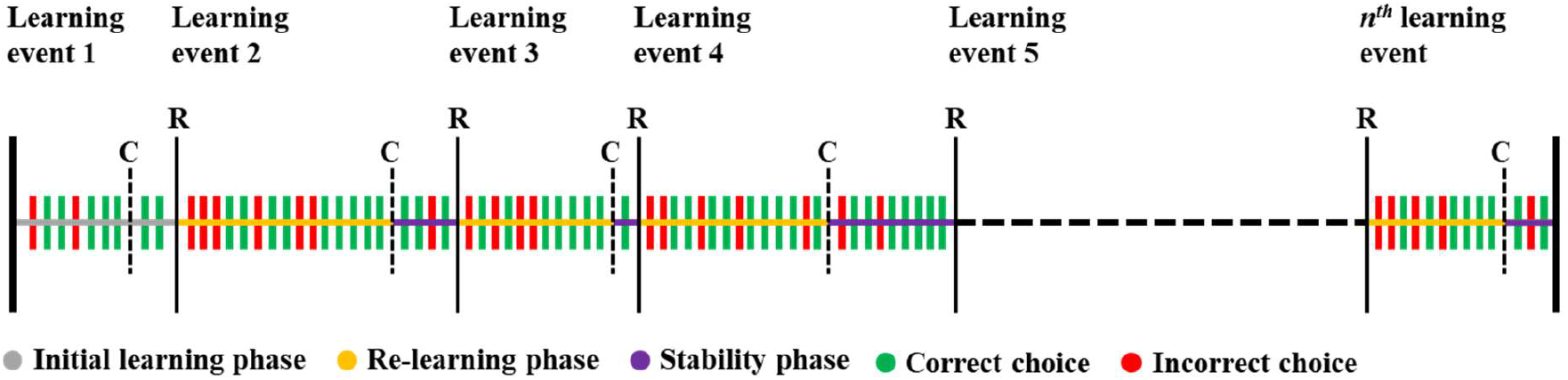
Trial and task phase overview of the serial reversal learning task. Dashed vertical lines show when criterion was reached (C); thin vertical lines show where outcome contingencies reversed (R) and a new learning event starts. Initial learning is the first learning event. After each reversal (R) participants are in the re-learning phase until they reach criterion (C). Participants are then in the stability phase until outcome contingencies reverse (R). The learning criterion must be maintained during the stability phase before reward contingencies reverse. Incorrect choices during the re-learning phase are defined as reversal errors, and the last reversal error of each re-learning phase is defined as the final reversal error. Each participant completes a total of 360 trials.

**Figure 3.**
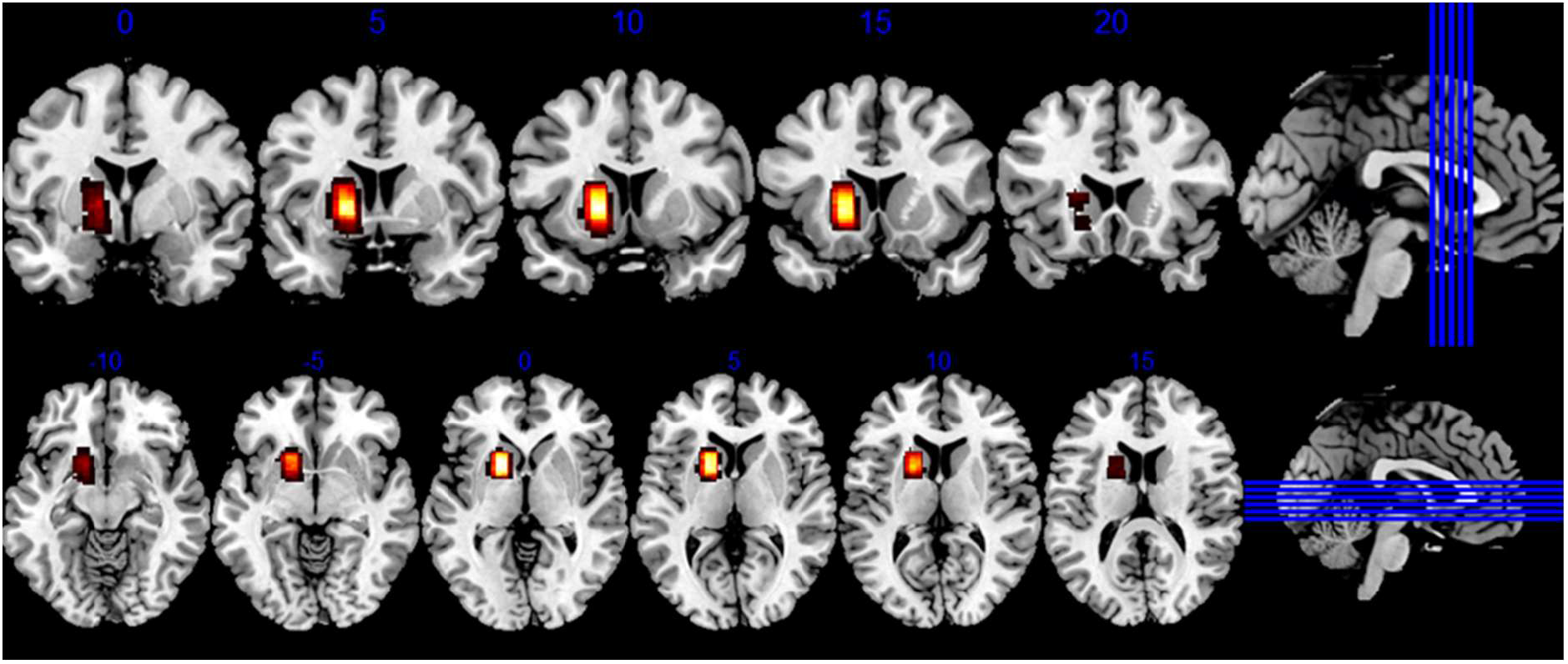
Coronal and axial slices visualising voxel positioning within the striatum for all participants in standard space. Heatmap denotes extent of spatial overlap, from maximum/yellow to minimum/red.

**Figure 4.**
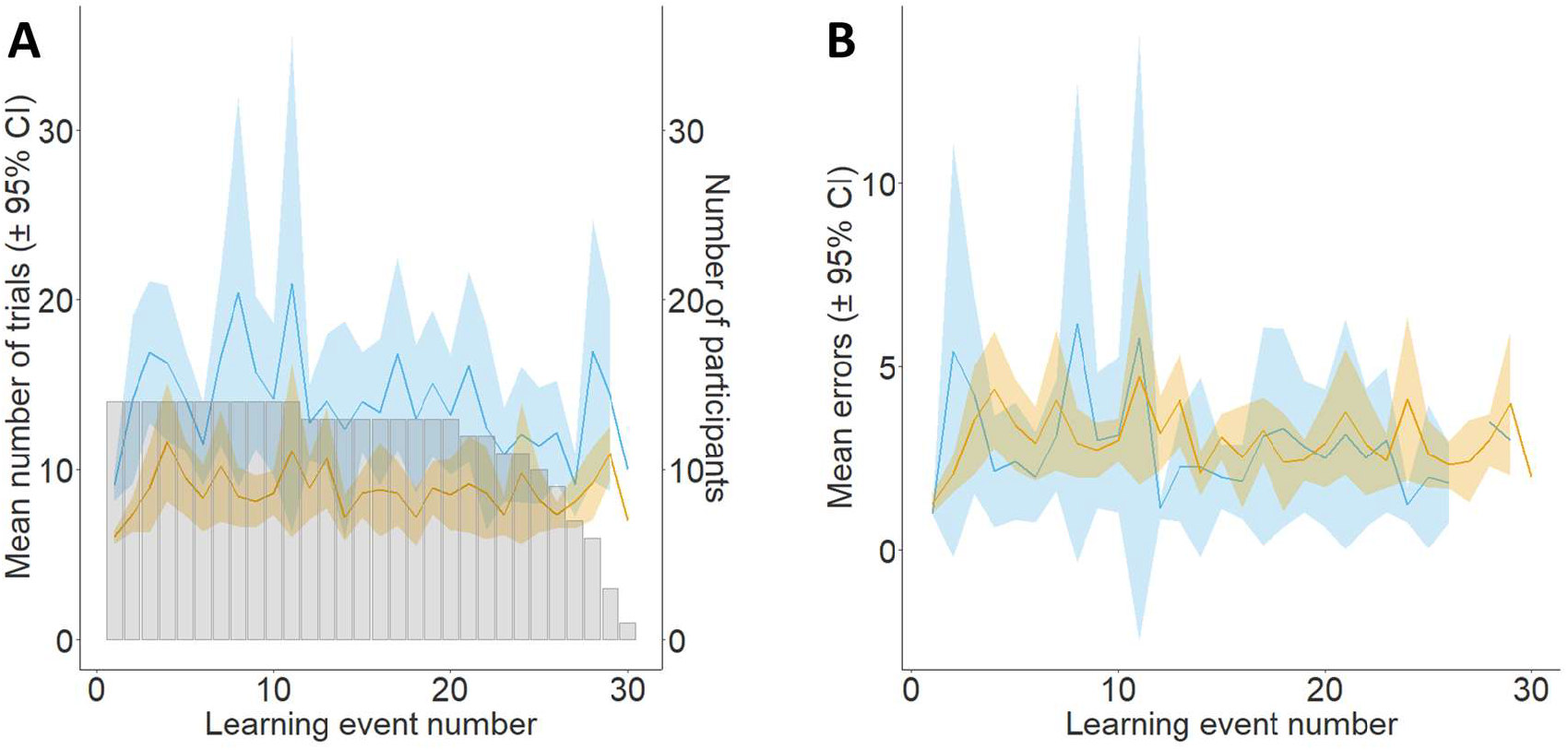
A: Average number of trials to reach criterion (orange) and reversal (blue) for each learning event (± 95% confidence intervals). Number of participants who reached each learning event (grey bars). B: Average number of perseverative (orange) and regressive (blue) errors for each learning event (± 95% confidence intervals).

**Figure 5.**
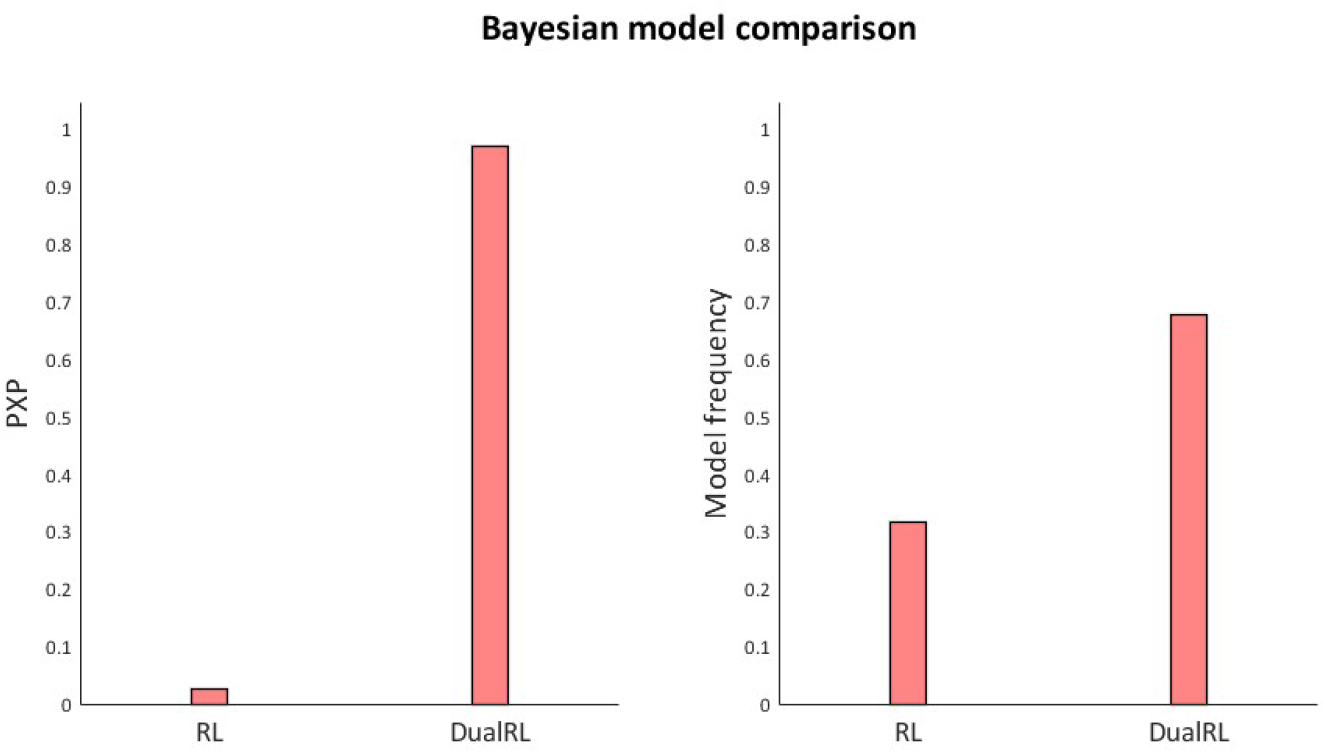
Protected exceedance probability and model frequency for the single and dual learning rate reinforcement learning models. The exceedance probability is the probability that a given model is the most commonly expressed model across particiapants, given the null hypothesis none of the models are sufficiently supported by the data

**Figure 6.**
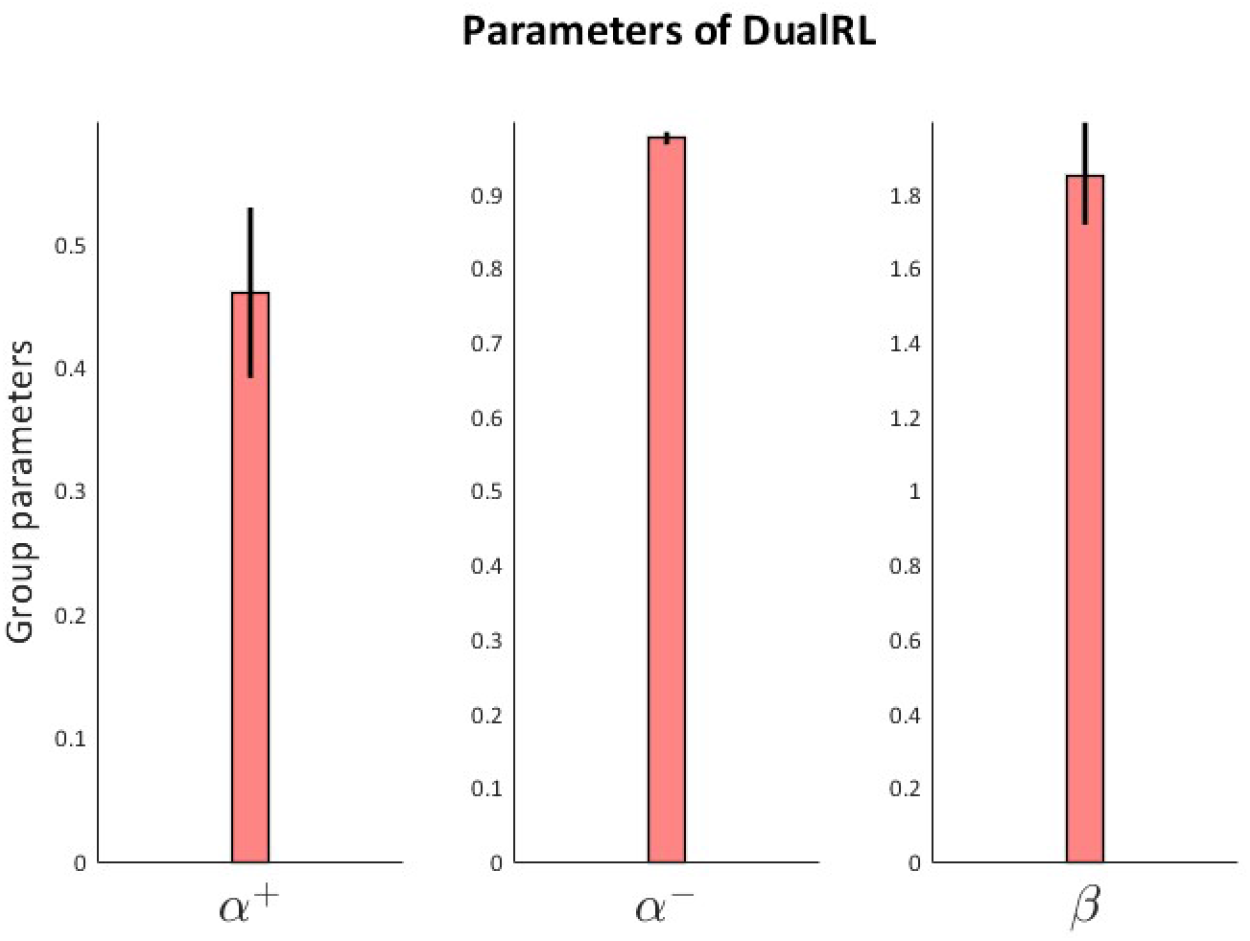
Group level parameter estimates for the learning rate for positive (α+) and negative (α-) prediction errors and the inverse temperature parameter (β). The error bars for all plots are the standard error of the mean.

**Figure 7.**
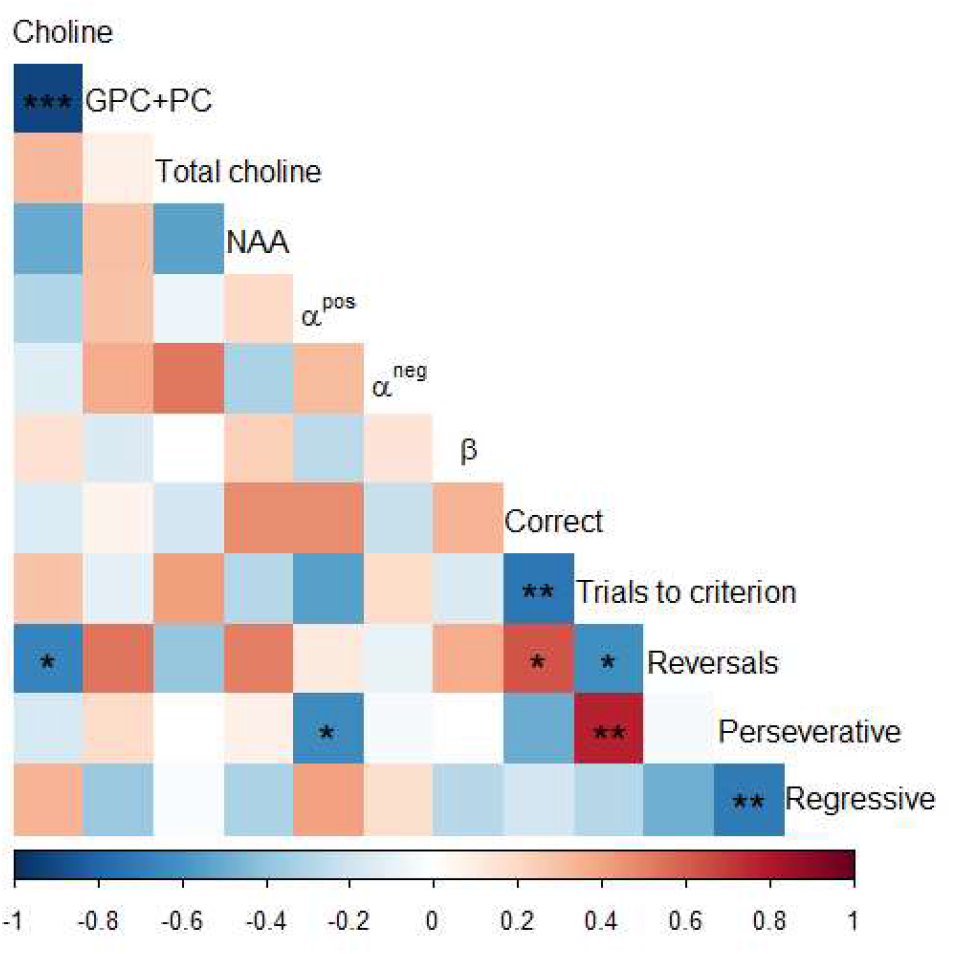
Correlations between metabolite concentrations, reinforcement learning model parameter estimates, and measures of reversal learning behaviour. Significant correlations are denoted with asterisks (*p < 0.05, **p < 0.01, ***p < 0.001)

At the beginning of the task, one of the two images were randomly assigned as the correct image, and the other as the incorrect image. The probability of winning points on the correct image was 0.8, and the probability of losing points was 0.2. The inverse was true for the incorrect image. Outcomes were pseudo-randomised such that the assigned probabilities were true for blocks of 20 consecutive selections of the correct or incorrect choice. Additionally, no more than six of the same outcomes (win or loss) would be consecutively presented for the correct or incorrect choice. If participants won, their cumulative total increased by 50. If they lost, their cumulative total decreased by 50. If they did not choose an image, their cumulative total did not change. For outcome probabilities to reverse participants had to reach and maintain a predefined learning criterion: the selection of the correct image on five of the previous six trials. After reaching criterion participants entered a stability phase where the probability of reversal was equal to the number of trials where criterion had been maintained, divided by 10 (adapted from Hampton et al. (2006)). If criterion was not maintained, then the probability of reversal was reset to 0 and restarted once criterion was reached. The reversal event involved the switching of outcome probabilities, with the correct image becoming incorrect and *vice versa*. After reversal, participants had to re-reach and maintain the learning criterion for the reassigned outcome probabilities before outcome probabilities would reverse again. Participants completed 360 trials of the reversal learning task. Participants completed 20 practice trials. Practice trials followed the same structure as trials in the scanner, but participants did not receive any feedback for their choices. Instead, hashtags were presented in place of outcome and cumulative total feedback.

### Computational Modelling

#### Overview

Two models were fit using mean-field variational Bayes to perform hierarchical Bayesian inference on our behavioural data using the MATLAB computational/behavioural modelling toolbox (Piray, Dezfouli, et al., 2019). Calculating parameter distributions at the population level using hierarchical Bayesian inference is advantageous over hierarchical parameter estimation, since one of the assumptions of hierarchical parameter estimation is that a given model is responsible for generating data from all subjects. During hierarchical parameter estimation, each participant equally influences group level parameters as model identity is included as a fixed effect despite it not necessarily being true that one given model best explains the behaviour of each subject. By contrast, hierarchical Bayesian inference takes a random effects approach to parameter estimation and model comparison (Piray & Daw, 2020).Therefore, an advantage of hierarchical Bayesian inference over hierarchical parameter estimation is that it includes a step where the responsibility of each model for generating a given dataset is calculated, and this responsibility influences group parameter estimation. The modelling approach described below follows recommendations from Wilson & Collins (2019).

#### Models

Model one is a model-free reinforcement learning model with a single learning rate parameter (α), and an inverse temperature parameter (β). The learning rate parameter defines the rate that value estimates are updated based on the difference between expected and actual outcomes, also known as a prediction error. The inverse temperature parameter describes the degree to which choices are based on value estimates. The lower an agent’s inverse temperature parameter, the more stochastic their choices will be. When β=0, choices would be made completely at random; when β=∞ the choice with the largest expected value would be deterministically chosen. In this model the softmax function is used to calculate the probability of making choice *k* at time 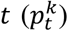, and is based on expected values *Q* and the inverse temperature parameter β. These probabilities are used during model fitting to calculate parameters that best describe the data. The softmax choice rule is defined as:

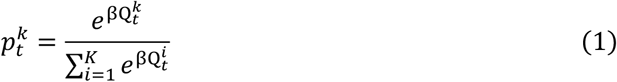

where 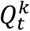 is defined as:

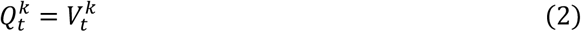

The expected value for choice *k* is updated such that 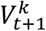 is equal to 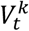 plus the product of the learning rate (α) and the prediction error (*δ*) (eq. 3). The prediction error *δ* is defined as the difference between the actual *λ*_t_ and expected value 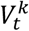 for choice *k* at time *t* (eq. 4).

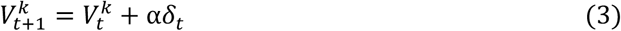

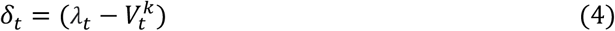

Model two is a model-free reinforcement learning model with separate learning rates for positive (α^+^) and negative (α^-^) prediction and errors (eq. 5) and an inverse temperature parameter (β). The softmax function (eq. 1) is used to calculate the probability of making choice *k* at time *t* 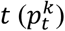. The expected value for choice *k* is updated such that 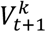 is equal to 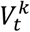 plus the product of the learning rate (α^+/-^) and the prediction error (*δ*) (eq. 5). The prediction error *δ* is defined as the difference between the actual *λ*_t_and expected value 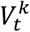 for choice *k* at time *t* (eq. 4). Separate learning rates for positive and negative prediction errors were included in this model because they have been shown to have asymmetric effects on expected value updating (Niv et al., 2012).

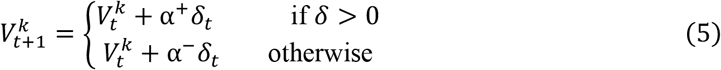

#### Model fitting

Model fitting and parameter estimation was performed using hierarchical Bayesian inference as implemented in the Computational/Behavioural Modelling toolbox (Piray, Dezfouli, et al., 2019). Model fitting was performed using behavioural data from the twenty-nine participants who successfully completed the reversal learning task; data from these participants were modelled to produce better estimates of group-level parameter distributions. In the first step of model fitting each model was fitted to participant’s data using Laplace approximation for non-hierarchical inference to generate a maximum a-posteriori estimates for each parameter for each subject, and log-model evidence for each subject. This non-hierarchical model fit requires that parameters have Gaussian priors; for all parameters these priors were specified as having mean=0, variance =6.25, in line with previous reports (Piray, Dezfouli, et al., 2019; Piray, Ly, et al., 2019). These values were selected because it creates a wide range of values that parameters could take. These values were then used during hierarchical Bayesian inference, implemented using mean-field variational Bayes. Each iteration of model fitting contained the following steps 1. Calculate summary statistics, 2. Update estimates of the posterior distribution for group parameters, 3. Update estimates of the posterior for individual parameters, 4. Update estimates of responsibility for each model in generating given data. Model fitting was iterated until the model reached convergence. The best fitting model was determined by the model with the highest exceedance probability, which is the probability that a given model is more commonly expressed than other candidate models in model space. Lastly, Model fitting was re-run, but this time under the hypothesis that observed differences in model fit are due to chance. This returns the protected exceedance probability, and a more conservative approach for finding the best fitting model (Piray, Dezfouli, et al., 2019).

### Magnetic Resonance Spectroscopy

#### Data acquisition

^1^H-MRS spectra and MR images were acquired at the Centre for Integrative Neuroscience and Neurodynamics, University of Reading, using a Siemens Magnetom Prisma-fit scanner 3T scanner and a 32 channel receiver head coil. High resolution T1-weighted anatomical images were acquired with a magnetization-prepared rapid gradient-echo (MP-RAGE) with GeneRalized Autocalibrating Partially Parallel Acquisitions (GRAPPA) (R = 2) sequence [TR = 2300ms; TE = 2.29ms; TI = 900ms slices = 192; voxel volume ≈ 0.9mm3; slice thickness = 0.94mm; distance factor = 50%; slice oversampling = 16.7%; FOV = 240× 240mm; matrix = 256 × 256; flip angle = 8°; phase encoding direction = A → P; echo spacing = 7ms]. T2 HASE images [TR = 1500 ms; TE = 82 ms; FOV = 220 × 220mm; flip angle = 150^°^; voxel = 0.7 × 0.7 × 3 mm; 15 slices] were acquired immediately prior to the acquisition of the MRS spectra; this positioned the axial plane of the voxel in the isocenter of the magnetic field, optimizing the homogenization of the magnetic field during shimming. The native scanner PRESS sequence [striatal voxel = 15 ×10 × 15 mm; TR = 2000 ms; TE = 30ms; 256 transients; water suppression bandwidth = 50 Hz; automatic shimming] was used to acquire spectra in the left dorsal striatum of all participants, similar to Bell et al. (2018), and Bell, Lindner, et al. (2019). The same PRESS sequence was used to acquire a water unsuppressed peak for eddy current correction and calculating absolute metabolite concentrations [15 transients]. PRESS data were acquired following the acquisition of echo planar data presented in Williams & Christakou, (2021).

#### Preprocessing

^1^H-MRS data analysis was performed in line with experts’ consensus recommendations published by (Near et al., 2020). MRS data pre-processing was carried out using the MATLAB toolbox FID-A (Simpson et al., 2015). Firstly, radiofrequency coil channels were combined, and bad individual spectra (spectra > 4 standard deviations used as rejection threshold) were removed. Spectra were then aligned to correct for frequency drift and averaged to create a single spectrum. The averaged spectrum was brought in phase (first-order phasing), and zero-order phase correction was applied using the creatine peak. The spectrum was frequency shifted so that creatine appeared at 3.027ppm for the water suppressed spectrum, and water appeared at 4.65ppm in the water unsuppressed spectrum. eddy current correction was applied to remove distortion in the spectrum due to fluctuations in the B0 field, then the water peak was subtracted from the water suppressed spectrum using Hankel-Lanczos Singular Value Decomposition (Pijnappel et al., 1992) as implemented in FID-A.

#### Spectral quantitation

A metabolite basis set was generated using the MATLAB FID-A toolbox (Simpson et al., 2015). Sixteen metabolites (acetate, aspartate, choline, creatine, gamma-aminobutyric acid (GABA), glucose, glutamate, glutamine, lactate, myo-inositol, N-acetyl aspartate (NAA), phosphocreatine, phosphocholine (PC), glycerophosphocholine (GPC), scyllo-inositol, and taurine) were simulated at a field strength of 3T using a PRESS pulse sequence (TE1 = 16.6 ms, TE2 = 13.4 ms, 4096 points, spectral width = 2399.8Hz, linewidth = 12.684Hz). Choline was modelled separately from PC and GPC, which were added following simulation to form a single peak (GPC+PC).

Automatic quantification of metabolites from the spectra were calculated using the jMRUI tool Accurate Quantification of Short Echo time domain Signals (AQSES) (jMRUI, version 6.0; http://www.jmrui.eu/; Garcia et al., 2010; Naressi et al., 2001; Stefan et al., 2009). The NAA peak in the spectra was shifted to 2.02 ppm to correct for chemical shift displacement, and the metabolite model was realigned with the NAA peak in the spectra. The following settings were used for quantification: equal phase for all metabolites; begin fixed timing; delta damping -10 to 40 Hz; delta frequency -10 to 10 Hz, no background handling; 0 truncated points; 4096 points in AQSES; normalization on. Metabolite concentrations were corrected by calculating their amplitude relative to the corresponding regional water peak (acquisition correction=1, tissue correction=0.5555).

MRS voxels were co-registered with high resolution T1 anatomical images using CoRegStandAlong in Gannet 3.1 and SPM-12 (Ashburner & Friston, 2005; Edden et al., 2014). During registration, the fraction of grey matter, white matter, and cerebrospinal fluid was calculated for each spectral acquisition. These fractional tissue compositions were used to correct the concentrations of choline and GPC+PC for partial volume and relaxation effects using the MATLAB toolbox MRSParVolCo (https://github.com/DrMichaelLindner/MRSParVolCo), based on the formulae described by (Gasparovic et al., 2006).

### Statistical analysis

Statistical analyses were performed using the R programming language (R Core Team, 2020; Wei & Simko, 2021; Wickham, 2016; Wickham et al., 2020) and SPSS (IBM Corp. Released 2017. IBM SPSS Statistics for Windows, Version 25.0. Armonk, NY: IBM Corp.). Correlational analyses were used to assess the relationship between metabolite concentrations, model parameter estimates, and behavioural performance. Hierarchical multiple regression was used to assess whether variance in choline concentrations could be explained by participants’ model parameter estimates and behaviour. As part of this analysis we included GPC+PC concentration, since we know it is anti-correlated with choline concentrations (Bell, Lindner, et al., 2019; Lindner et al., 2017), and number of reversals as covariates of no interest. Lastly, to assess the specificity of these results to choline, we re-ran our regression analysis using NAA.

## Results

### Reversal learning performance

Participants made correct choices at significantly greater than chance level (mean correct choices = 252.71, 95% CI [244, *∞*], *t*(13) = 14.19, *p* < .001, SD = 19.18, Range = 193 - 269), and experienced an average of 24.21 (SD = 4.96; Range = 10-29) reversals. The average number of trials taken to reach criterion was 8.756 (SD = 4.371; Range = 5 - 40); an average of 3.044 (SD = 2.161; Range = 0 - 17) perseverative errors were made following the reversal of contingencies before reaching criterion in each learning event. On 3.357 (SD = 3.734; Range = 0 - 12) trials participants did not respond to either of the two images presented to them. An average of 4617.857 (SD = 1099.407, Range = 1200 - 5550) points were collected by the end of the task. The average time taken by participants to make a choice following the onset of the stimuli was 595.746 milliseconds (SD = 110.104, Range = 398.713 - 782.612).

### Model fit

Two reinforcement learning models were fit to participants behavioural data from the reversal learning task. The first was a model-free reinforcement learning model with a single learning rate α, and an inverse temperature parameter β. The second model was a model-free reinforcement learning model with two learning rates, α^+^ and α^-^ for positive and negative prediction errors, and an inverse temperature parameter β. Model fitting was performed using hierarchical Bayesian inference with the MATLAB Computational/Behavioural Modelling toolbox (Piray, Dezfouli, et al., 2019). Overall, the dual-learning rate model out-performed the single learning rate model with respect to its protected exceedance probability (0.9727 for the dual learning rate model, 0.0273 for the single learning rate model), and the goodness of fit to each participant’s data (the dual learning rate model had a higher responsibility for eleven of the fourteen participants included in the analysis of spectroscopy data). The group-mean learning rate for positive prediction errors (α^+^) was 0.4605 and 0.9786 for negative prediction errors (α^-^); the group-mean inverse temperature parameter (β) was 1.8533.

### Metabolite quantitation

Metabolite spectra were quantified for twenty-one participants to measure concentrations of choline and glycerophosphocholine plus phosphocholine (GPC+PC) in the dorsal striatum. Separate measures of choline and GPC+PC could be quantified for fourteen participants. The mean concentration of choline in the dorsal striatum was 0.797 millimolar (mM) (SD = 0.253, Range = 0.481 - 1.351); for GPC+PC the mean concentration was 0.854 mM (SD = 0.241, Range = 0.323 - 1.238). As previously reported, we found that concentrations of choline and GPC+PC were anti-correlated (*r* = -0.912 *t*(11) = -7.379, 95% CI = [-0.974, -0.726], *p* < 0.001) (Bell et al., 2018; Lindner et al., 2017; Miller et al., 1996).

### Relationship between choline measures and behaviour

To investigate the relationship between neurochemistry and behaviour we first ran correlations between our measures of task performance (correct choices, perseverative errors, regressive errors, and number of reversals), modelling parameters (α^+^, α^-^, β) and metabolite concentrations in the dorsal striatum (Cho, GPC+PC, total choline, and NAA). Increases in dorsal striatal choline were associated with decreases in dorsal striatal GPC+PC (*r* = -0.912 *t*(11) = -7.379, 95% CI = [-0.974, -0.726], *p* < 0.001), and decreases in the number of reversals participants experienced (*r* = -0.667 *t*(11) = -2.971, 95% CI = [-0.891, -0.184], *p* = 0.013). Increases in the number of reversals was associated with an increase in the number of correct responses made (*r* = 0.630 *t*(11) = 2.688, 95% CI = [0.120, 0.877], *p* = 0.021). An increase in the number of perseverative errors was associated with a decrease in the learning rate for positive prediction errors (α^+^; *r* = -0.632 *t*(11) = -2.701, 95% CI = [-0.877, -0.123], *p* = 0.021), and a decrease in the number of regressive errors made (*r* = - 0.709 *t*(11) = -3.333, 95% CI = [-0.906, -0.259], *p* = 0.007). No other significant correlations were identified at a threshold of *p* < 0.05.

Before running our hierarchical multiple regression, the following assumptions were tested. We tested for multicollinearity by calculating variance inflation factors (VIF) for our predictor variables; no variables had a VIF greater than 10, suggesting they have not violated assumptions of multicollinearity (Field, 2013). To test the assumption that the residuals of predictors are uncorrelated, we used the Durbin-Watson test and found this assumption was met (Durbin-Watson = 2.288); values of two suggest residuals are uncorrelated, while values less than one or greater than three suggest indicate there are problematic positive or negative correlations (Field, 2013). Plots of standardised residuals and standardised predicted values suggested the assumptions of homogeneity of variance were met; P-P plots of standardised residuals suggested that the assumption of normality may have been violated, however regression results are unlikely to be biased by violations of normality when there are ten or more observations for each variable (Schmidt & Finan, 2018). Cook’s distance for two participants were greater than one and suggests these participants may have a disproportionate influence on the model, however these participants were not excluded from analysis given the size of the dataset.

A two-stage hierarchical multiple regression model was run to investigate the relationship between levels of choline and task performance. The GPC+PC concentration and number of reversals were included in the first stage as a covariates of no interest. Perseverative and regressive errors, positive (α^+^) and negative (α^-^) learning rates, and the inverse temperature parameter (β) were included in the second stage of the model. One participant was excluded from our regression model because their behaviour suggested they did not understand the task, but this did not influence model fitting. The first stage of the regression revealed that GPC+PC and the number of reversals were significant predictors of choline concentrations in the dorsal striatum *F* (2,12) = 35.522, *p* < 0.001, and explained 87.6% of the variance in choline concentration. The inclusion of model parameters and number of errors at the second stage of the regression increased the variance explained by 10.6%, and this was a significant increase in explained variance *F* (7,12) = 39.802, *p* < 0.001, *f*^*2*^ = 0.119 (Table 1).

**Table 1.** Hierarchical regression model predicting concentrations of choline. At the first stage GPC+PC concentration and the number of reversals are included as covariates of no interest. In the second stage reinforcement learning model parameter estimates, perseverative and regressive errors are included in the model.

| Variable | Unstandardised $\beta$<br>/ Standardised $\beta$ | $t$ | R | $R^2$ | $\Delta R^2$ |
| --- | --- | --- | --- | --- | --- |
| Stage 1 |  |  | .936 | .876 |  |
| GPC+PC | -.820 / -.778 | -5.913*** |  |  |  |
| Reversals | -.023 / -.250 | -1.903 |  |  |  |
| Stage 2 |  |  | .991 | .982 | .149* |
| GPC+PC | -.894 / -.848 | -7.083** |  |  |  |
| Reversals | -.037 / -.400 | -3.407* |  |  |  |
| $\alpha^+$ | -.493 / -.303 | -2.914* | | | |
| $\alpha^-$ | 1.891 / .328 | 3.275* | | | |
| $\beta$ | -.067 / -.125 | -1.192 | | | |
| Perseverative err | -.014 / -.619 | -4.397** |  |  |  |
| Regressive err | -.015 / -.574 | -4.197** |  |  |  |

$N = 13$ ; \* $p < .05$ ; \*\* $p < .01$ ; \*\*\* $p < .001$

**Table 2.** Hierarchical regression model predicting concentrations of NAA. NAA model was used as a control metabolite to demonstrate the specificity of the model for choline. At the first stage GPC+PC concentration and the number of reversals are included as covariates of no interest. In the second stage reinforcement learning model parameter estimates, perseverative and regressive errors are included in the model.

| Variable | Unstandardised $\beta$<br>/ Standardised $\beta$ | $t$ | R | $R^2$ | $\Delta R^2$ |
| --- | --- | --- | --- | --- | --- |
| Stage 1 |  |  | .504 | .254 |  |
| GPC+PC | .091 / .033 | .103 |  |  |  |
| Reversals | .117 / .485 | 1.501 |  |  |  |
| Stage 2 |  |  | .889 | .790 | .536 |
| GPC+PC | 1.836 / .672 | 1.628 |  |  |  |
| Reversals | -.018 / -.073 | -.181 |  |  |  |
| $\alpha^+$ | 4.083 / .967 | 2.701* | | | |
| $\alpha^-$ | -16.326 / -1.092 | -3.164* | | | |
| $\beta$ | 1.367 / .976 | 2.703* | | | |
| Perseverative err | .059 / .995 | 2.050 |  |  |  |
| Regressive err | .045 / .668 | 1.417 |  |  |  |

$N = 13$ ; \* $p < .05$ ; \*\* $p < .01$ ; \*\*\* $p < .001$

To test whether these results were specific to the concentration of choline, we re-ran the same regression model with the concentration of N-acetyl aspartate (NAA) as our dependent variable. GPC+PC and the number of reversals were not significant predictors of NAA concentration in the first stage of the regression model *F* (2,12) = 1.704, *p* = 0.231, and explained 25.4% of the variance in NAA concentration. The addition of our modelling parameters and measures of error increased the explained variance in NAA to 79%, however this was not a significant increase in explained variance and the model did not significantly explain concentrations of NAA *F* (7,12) = 2.693, *p* = 0.146.

## Discussion

We used ^1^H-MRS to investigate whether variability in resting levels of choline in the dorsal striatum are related to reversal learning performance and reinforcement learning model parameter estimates in a two-alternative serial reversal learning task. Positive and negative learning rates, and the number of perseverative and regressive errors were significant predictors of dorsal striatal choline, and their inclusion in our regression model explained significantly more variance in choline than when we only included GPC+PC and number of reversals as covariates of no interest. Almost all the variability in choline was explained by this regression model (98.2%). Conversely, when we re-ran the same analyses for a metabolite that we consider a functional control we observed no association with performance. Specifically, using NAA as our predicted outcome, we found that neither level of our hierarchical model significantly predicted NAA concentrations.

Further correlational analyses are in line with previous findings. Firstly, we show that levels of choline in the dorsal striatum are inversely correlated with GPC+PC concentrations (Bell, Lindner, et al., 2019; Lindner et al., 2017). We believe this relationship is not due to issues with model fitting, as Lindner et al. (2017) previously demonstrated using synthetic spectra that relative differences in concentrations of choline and GPC+PC are faithfully recovered during model fitting when separate peaks for choline and GPC+PC are used. Secondly, we found that concentrations of choline are rest are negatively correlated with the number of reversals participants made. This mirrors previous findings that participants who were quicker to reverse had lower levels of dorsal striatal choline at rest (Bell, Lindner, et al., 2019), and that participants who learned during reversal had lower levels of choline at rest than those who did not (Bell et al., 2018).

Previous work on deterministic reversal learning by D’Cruz et al. (2011) has demonstrated that unexpected outcomes during two- and four-choice tasks activate similar brain areas, but that activation during four choice learning is significantly greater in several regions, including the thalamus. Although D’Cruz et al. (2011) do not localise this activation to specific nuclei in the thalamus, based on our understanding of the roles of thalamostriatal connections in flexibility, especially in signalling unexpected outcomes (Bell, Langdon, et al., 2019; Bradfield et al., 2013; Matsumoto et al., 2001), we would expect this cluster to include the centromedian-parafascicular nuclei. If the centromedian-parafascicular nuclei show different levels of activation in two and four choice reversal learning, then their efferent projections to the cholinergic interneurons of the dorsal striatum may have different effects on the striatal cholinergic system during reversal learning. If resting levels of choline were consistent for participants completing both reversal learning tasks, then variability in input to the striatal cholinergic system could have a differential effect on subsequent behaviour in the two tasks.

Here we provide evidence suggesting choline levels at rest may have dissociable effects on behaviour, depending on task context. During multi-alternative reversal learning, there is a protracted period of learning where the participant’s experience closely mimics the experience in animal studies, with no prior knowledge of task context or structure. Conversely, in serial reversal learning, such as the task used here, participants are provided with instruction about the general structure of the task which can be used to scaffold task representations. This prior knowledge should enable participants to readily form mature task representations. Once participants develop an “ if not A, then B” heuristic for choice, their task representation can be considered as “ saturated”, as no other contextual information is available that might further support adaptive behaviour. However, task representations in the multi-alternative task can be considered as “ unsaturated”, because participants only form mature task representations after experiencing both the protracted periods of initial and reversal learning. These differences could explain why resting cholinergic “ tone” has dissociable effects during serial and multi-alternative reversal learning. Here, we found that dorsal striatal choline concentration was negatively associated with perseverative errors but positively associated with the learning rate for negative prediction errors. Conversely, Bell, Lindner et al. (2019) found that dorsal striatal choline was positively correlated with perseverative errors, but negatively correlated with the learning rate for negative prediction errors. In an unsaturated context (multi-alternative task), a low cholinergic tone may be beneficial for generating contrast between periods of stability and change. This increased contrast could detect change more clearly, enabling flexibility when required. However, in the saturated context (serial reversal learning task), a high cholinergic tone may be more beneficial: participants with higher choline at rest were influenced more by negative than positive prediction errors and showed fewer perseverative and regressive errors. By contrast, lower cholinergic tone may disproportionately increase learning from positive feedback for perseverative errors and decrease learning from negative feedback from regressive errors. These participants would then be slower to reverse and less likely to maintain behaviour. However, these participants do not appear to be any more stochastic in their behaviour than participants with higher cholinergic tone, as the inverse temperature parameter did not significantly predict choline levels at rest. Task performance in different contexts thus appears to be modulated by the state of the cholinergic system at rest, with the saturation of task representation modulating its relationship with performance.

One potential factor that may also account for differences between the serial and multi-alternative reversal learning tasks is the model fitting procedure. Although both studies use the same reinforcement learning model, Bell, Lindner et al. (2019) fitted their model to initial and reversal learning separately, while here we fit the model to all trials at once. Therefore, parameter estimates in both studies do not describe exactly equivalent aspects of behaviour. This difference is inevitable due to the structure of our task. Combining trials across different phases would produce invalid results, as the model would be fit to non-contiguous trials that do not reflect the experience of the participant. Additionally, the model fitting approach taken here uses Hierarchical Bayesian Inference, while Bell, Lindner et al. (2019) use Maximum Likelihood Estimation. Hierarchical Bayesian methods have been shown to more accurately provide point estimates of individual parameters (Farrell & Ludwig, 2008; Katahira, 2016), and therefore some difference between our results and those of Bell, Lindner et al. (2019) may also be due to model fitting approaches. Most probable is that differences between our results and those of Bell, Lindner et al. (2019) are due to a combination of psychological and methodological factors, given none of the explanations given are mutually exclusive of one-another.

Given the preliminary nature of this study, it is important that the results presented here are received with caution. Despite strong evidence from the animal literature and from our previous work in humans describing the roles of the striatal cholinergic system in flexibility (e.g. Bell et al., 2018; Bell, Lindner, et al., 2019; Bradfield et al., 2013; Brown et al., 2010; Ragozzino et al., 2002, 2009), these results need to be replicated. Several limitations of this study are related to the sample size. Firstly, although data were acquired for thirty-three participants, only thirteen datasets were used in our final analysis. Of the twenty excluded participants, thirteen had issues related to the acquisition or analysis of their metabolite spectra. One potential explanation for this data loss is the ^1^H-MRS spectra were acquired following the acquisition of fMRI data using echo-planar imaging sequences, which can cause frequency drift, leading to the distortion of metabolite spectra (El-Sharkawy et al., 2006; Harris et al., 2014; Lange et al., 2011). Future work could aim to minimise the effects of frequency drift by leaving time for the gradient coils to cool prior to the acquisition of spectroscopy data. Further, given the size of the sample size reported here it is worth noting that the implied α for ΔR^2^ (based on *f*^2^, sample size, and number of predictors) in the stage 2 choline regression model was 0.183. This inflated α increases the risk of type 1 error, so the results reported here may have an increased likelihood of being due to type 1 error. Furthermore, two participants had Cook’s distances that were greater than the recommended threshold and therefore these participants could disproportionately affect regression coefficient estimation (Field, 2013).

As it stands, and in line with previous work, this study provides further evidence for the role of the dorsal striatal cholinergic system in flexibility, and particularly during reversal learning. We find that levels of choline are associated with learning rates for positive and negative prediction errors, and the number of perseverative and regressive errors. These findings show that the dorsal striatal cholinergic system appears to be involved in producing flexible behaviour during reversal learning, in line with previous work. Importantly, however, we find potential differences in how the system may be involved during instructed serial reversal learning and during uninstructed multi-alternative probabilistic reversal learning. Together, these results suggest an important role for the cholinergic system in flexibility in humans, as is suggested by findings from animal literature, and describe how the system may function in different contexts.

## Acknowledgements

The authors would like to thank Tiffany Bell for her advice on the spectral quantitation of choline containing compounds.

